# Exogenous Melatonin accelerates seed germination in cotton(*Gossypium hirsutum* L.)

**DOI:** 10.1101/618959

**Authors:** Shuang Xiao, Liantao Liu, Hao Wang, Dongxiao Li, Zhiying Bai, Yongjiang Zhang, Hongchun Sun, Ke Zhang, Cundong Li

**Affiliations:** Key Laboratory of Crop Growth Regulation of HeBei Province, Hebei Agricultural University, Baoding, Hebei Province, China

## Abstract

Seed germination is considered as the beginning of the spermatophyte lifecycle, as it is a crucial stage in determining subsequent plant growth and development. Although many previous studies have found that melatonin can promote seed germination, the role of melatonin in cotton germination remains unexamined. The main objective of this study is the characterization of potential promotional effects of melatonin (at doses of 0, 10, 20, 50, 100 and 200 μM) on cotton seed germination. This experiment demonstrated that low concentrations of melatonin can promote germination, while high concentrations failed to promote germination and even inhibited germination. Together, these results indicate that a 20 μM melatonin treatment optimally promotes cotton seed germination. Compared with the control, germination potential (GP), germination rate (GR) and final fresh weight (FW) increased by 16.67%, 12.30% and 4.81% respectively. Although low concentrations of melatonin showed some improvement in vigor index (VI), germination index (GI) and mean germination time (MGT), these effects did not reach significant levels. Antioxidant enzyme activity during seed germination was most prominent under the 20 μM melatonin treatment. Superoxide dismutase (SOD) and peroxidase (POD) activities were significantly increased by 10.37–59.73% and 17.79–47.68% compared to the melatonin-free control. Malondialdehyde (MDA) content was reduced by 16.73–40.33%. Two important plant hormones in seed germination were also studied. As melatonin concentration increased, ABA content in seeds decreased first and then increased, and GA_3_ content showed a diametrically opposite trend, in which the 20 μM melatonin treatment was optimal. The 20 μM melatonin treatment reduced ABA content in seeds by 42.13–51.68%, while the 20 μM melatonin treatment increased GA_3_ content in seeds to about 1.7–2.5 times that of seeds germinated without melatonin. This study provides new evidence suggesting that low concentrations of melatonin can promote cotton seed germination by increasing the activity of antioxidant enzymes, thereby reducing the accumulation of MDA and regulating plant hormones. This has clear applications for improving the germination rate of cotton seeds using melatonin.

## Introduction

Seed germination, an early and crucial stage in the spermatophyte life cycle, refers to the physiological process starting from the uptake of water by the dry seed and ending with radicle protrusion[1]. It is a complex process regulated by many coordinated metabolic, cellular and molecular events in the life cycle of higher plants.

Cotton (*Gossypium hirsutum* L.) is a globally important cash crop, fiber crop and oil crop, and it plays an important role in China’s economic development. Although cotton seeds have a long storage life, they often emerge poorly in early spring because of low seed vigor. For seeds harvested in wet years, the germination rate is often even lower, resulting in weak emergence after sowing and ultimately reducing cotton yields. Therefore, the quality of seed germination is related to the growth of subsequent cotton seedlings and resulting cotton yields.

Melatonin (*N*-acetyl-5-methoxytryptamine), a highly conserved and efficient molecule, exists widely in plants and animals[2]. The tryptophan-derived compound is synthesized in the hypothalamus[3] and was first isolated from the pineal gland of cattle in 1958, suggesting it only existed in animals. However, since melatonin was first detected in nine edible plants, including tomatoes (*Lycopersicon esculentum*) and cucumbers (*Cucumis sativus*) in 1995[4], melatonin has been detected in more than 300 plants, including some in monocotyledonous and dicotyledonous families[5–7], and the biosynthetic pathways and physiological functions of melatonin in plants have been extensively studied[8]. Melatonin has been detected in the seeds, roots, fruits and leaves of plants[9, 10]. In recent years, many studies have shown that melatonin in plants is synthesized from tryptophan and plays an important role in regulating plant growth, development and coping with various environmental stresses[10–12], including the through the regulation of seed germination[13]. Moreover, the effect of melatonin on seed germination differs in a concentration-dependent manner such that higher concentrations of melatonin inhibit or do not affect seed germination, while lower concentrations promote seed germination[14–16].

The production of reactive oxygen species (ROS) during seed aerobic metabolism leads to lipid peroxidation, which is an important cause of seed deterioration and inhibition of seed germination. Melatonin is the most potent endogenous compound known to scavenge free radicals, at a rate twice as high as that of vitamin E and four times as high as glutathione[17, 18]. In normal life activities, plants produce high levels of ROS through enzymatic reactions and leakage of electron transport chains during photosynthesis and respiration. Under normal conditions, cells achieve homoeostasis between ROS production and removal through antioxidant systems, thus maintaining low ROS levels. Previous research has revealed that melatonin protects algae from oxidative damage caused by H_2_O_2_[19] and protects plants from stress caused by drought[20, 21], low temperature[22], heavy metals[23, 24], and ultraviolet radiation[25], among other stressors, effectively preventing oxidative damage to their cellular structures and biomacromolecules. As a reproductive organ, seeds are vulnerable to oxidative damage, but the activity of antioxidant enzymes in seeds is very low, while melatonin has the ability to enhance the activities of scavenging enzymes[26, 27], such as superoxide dismutase (SOD; EC 1.15.1.1) and peroxidase (POD; EC 1.11.1.7), which play a crucial role in decreasing and eliminating ROS. Ultimately, melatonin can protect seeds from environmental damage, ensuring the smooth reproduction of plants[28].

In 2004, Hernández-Ruiz et al. first proposed that melatonin may be a plant growth hormone[14]. It is well known that phytohormones can affect the dormancy and germination of seeds and are important signal molecules for transmitting environmental changes during seed germination. At the same time, they have been extensively studied in dicots and monocots[29]. Abscisic acid (ABA) and gibberellins (GAs) are considered to be the main hormones in seed germination[30]. The dynamic balance of synthesis and catabolism of ABA and GA is the key to complete germination of seeds. Gibberellins, a group of important plant hormones, were first isolated from the pathogenic fungus *Gibberella fujikuroi* in 1938 by Japanese scholars[31]. GAs regulate the overall growth and development of plants and participate in controlling various plant development processes. At present, more than one hundred kinds of gibberellins have been found, with only a few, e.g., GA_1_ and GA_3_, demonstrating activity. Most related compounds are gibberellin synthesis precursors or products inactivated by metabolism. The active components of plant gibberellins mainly consist of GA_1_, GA_3_, GA_4_ and GA_7_. GAs can break seed dormancy, promote seed germination, trigger seed germination in suitable environments and time periods and prepare for subsequent seedling growth[32]. ABA is also an important hormone in the regulation of plant growth and development and accordingly plays a critical role in regulating various physiological processes in plants. It is derived from carotenoid precursors, which can delay seed germination and promote seed dormancy. ABA levels increase when plants are subjected to certain abiotic stresses, such as water deficit, and during seed development.

The positive effect of melatonin on seed germination observed in several plant species motivated this research. The objectives of the study were to investigate (1) whether melatonin can improve germination of cotton seeds, (2) whether melatonin can improve the physiological activity of seeds and (3) whether melatonin affect the hormones that regulate seed germination. This study aims to provide insight into the roles of melatonin in the germination of cotton seeds.

## Materials and methods

### Regents

Melatonin (*N*-acetyl-5-methoxytryptamine) was obtained from Sigma-Aldrich (St. Louis, MO, USA). All other reagents were obtained from Sinopharm Chemical Reagent Beijing Co., Ltd, China. All chemicals used in all experiments were of analytical grade.

### Plant material

The experiment was conducted in the Yellow River basin at Hebei Agricultural University, Baoding (38.85°N, 115.30°E), Hebei Province, China. ‘NDM601,’ a high-yielding commercial transgenic cotton (*Gossypium hirsutum* L.) cultivar, was used in the experiment.

### Germination tests

Selected cotton seeds were surface sterilized with 1% NaClO for 30 min and washed extensively with deionized water. Fifty seeds were then placed in Petri dishes (15 × 15 cm) containing double filter paper (Whatman International Ltd., Maidstone, UK) wet with 10 mL of the different treatment solutions and incubated in the dark at 25°C in an incubator. The following treatments were used: M0 (0 μM melatonin, distilled water only), M10 (10 μM melatonin), M20 (20 μM melatonin), M50 (50 μM melatonin), M100 (100 μM melatonin), M200 (200 μM melatonin). Germination tests were performed with six replicates. The solutions were renewed every 24 hours to maintain unaltered concentrations. At day 7 of the incubation, another batch of samples was harvested for the determinations of seedling fresh and dry weights and of seed germination rates. Germination assay samples were harvested at days 2, 4 and 6 of the incubation for biochemical and physiological measurements.

### Seed germination assessment

The germinated seeds (assessed when the emerging cotyledon was about the half-length of the seeds) were scored daily. Germination potential (GP) is the percentage of germinated seeds per Petri dish by day 3 of the germination assay. Germination rate (GR) refers to the percentage of germinated seeds per Petri dish by day 7. Germination index (GI) was calculated using the method developed by Wang et al.[33] as GI =∑(*G_i_/T_i_*), where *G_i_* is the germination percentage of the *i*^th^ day, and *T_i_* is the day of the germination test. High GI values indicate high seed quality and performance[34]. Vigor index (VI) = GI × fresh weight (FW) of germinated seeds on day 7. Mean germination time (MGT) was calculated according to the method by Ellis and Roberts[35], i.e., MGT = ∑(*N_i_T_i_*)/ ∑*N_i_*, where *N* is the number of seeds germinated at time *i*, and *T_i_* is day of the germination test. The lower the MGT value, the faster a population of seeds has germinated[36].

### Measurement of SOD, POD and MDA activities

Superoxide dismutase (SOD, EC1.15.1.1) activity was estimated based on the method by Giannopolittis and Ries[37] using the photochemical nitro blue tetrazolium chloride (NBT) method. Peroxidase (POD; EC 1.11.1.7) activity was measured at 25°C according to the method by Scebba et al.[38]. Malonaldehyde (MDA) content was determined following the method by Hodges et al. (1999)[39].

### Phytohormone quantification Determination of ABA and GA_3_

GA_3_ and ABA concentrations were determined using an indirect ELISA technique [40], the antibodies used were all monoclonal antibodies provided by China Agricultural University.

### Statistical analysis

Analysis of variance [41] was statistically performed using IBM SPSS Statistics 22.0 (IBM Corp., Armonk, NY, USA). All data are reported as mean ± standard deviation (SD) values. We performed six independent replicates for each treatment. The statistical significance was considered to be significant when the *P*-value was less than 0.05 in a one-way ANOVA.

## Results

### Germination rate, fresh weight, mean germination time and germination index

The response of plants to melatonin is highly concentration dependent [42]. We conducted an extensive set of germination assays using cotton seeds to determine how different concentrations of melatonin (0–200 μM) affect seed germination. The seeds started to germinate on the second day of incubation at 25°C. As Table 1 shows, GR increased first and then decreased as melatonin concentration increased. Among the treatments with different melatonin concentrations, the 20- μM melatonin treatment displayed the maximal effect on germination promotion. Compared with the control group (M0), the GR of the M10, M20 and M50 treatment groups increased by 10.40%, 12.30% and 1.46%, respectively, but did significantly differ from the control. When the melatonin concentration increased to 100 μM, the GR began to decrease. The GR and GP of the M100 treatment were significantly lower than those of the M0 group, by 12.59% and 21.67%, respectively. As the concentration of melatonin was further increased, the GR and GP significantly decreased. The GR and GP of the M200 treatment decreased by 32.21% and 36.48%, respectively. This indicates that low melatonin concentrations (M10–M50) promoted the GR and GP of cotton seeds, but after a critical concentration, melatonin had a serious inhibitory effect on the GR and GP of seeds. As the melatonin concentration increased, the inhibition of seed germination was enhanced further.

**Table 1.**
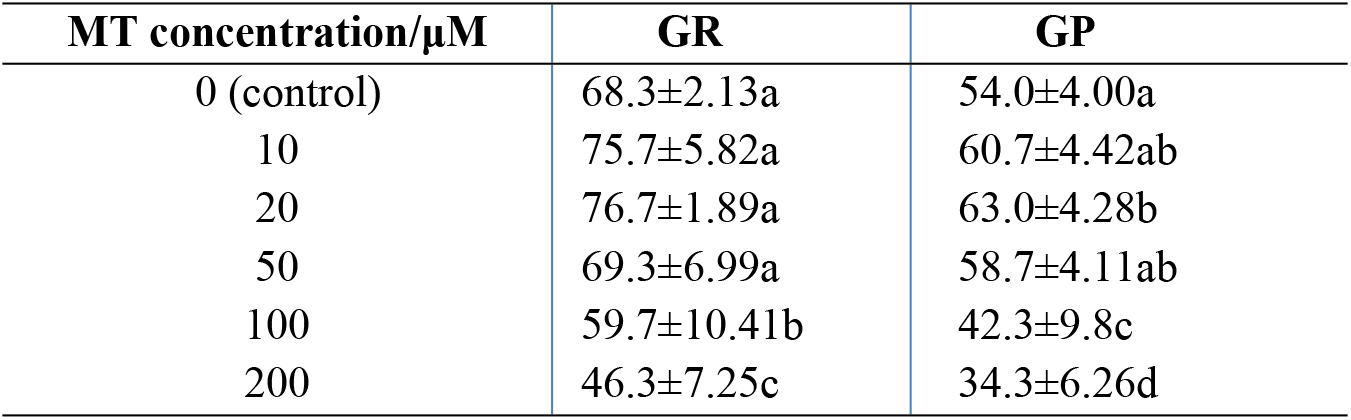
Effects of different melatonin concentrations on GR and GP of cotton seeds

| MT concentration/μM | GR | GP |
| --- | --- | --- |
| 0 (control) | 68.3±2.13a | 54.0±4.00a |
| 10 | 75.7±5.82a | 60.7±4.42ab |
| 20 | 76.7±1.89a | 63.0±4.28b |
| 50 | 69.3±6.99a | 58.7±4.11ab |
| 100 | 59.7±10.41b | 42.3±9.8c |
| 200 | 46.3±7.25c | 34.3±6.26d |

By focusing solely on the GP of the seeds, it is clear that the M20 treatment seeds germinated well. Under the M20 treatment, the GP of the seeds was 16.67% higher than that of the M0 group and reached a significant level. During seed germination days 4–7, the GR of the M20 treatment group continued to increase but not significantly. Therefore, this data indicates that the promotion of seed germination by melatonin is likely to occur at the early stage of seed germination. The GP refers to the germination speed and uniformity of seeds germinated, indicating the general vigor of the seed. Seeds with high GP values are highly active, and the resulting seedlings tend to grow uniformly and vigorously. Accordingly, VI is often used to evaluate seed quality. At day 7 of germination, VI and GI were significantly decreased (*P* < 0.05) while MGT was reduced in the M100 and M200 treatments. On day 7 of seed germination, we obtained the fresh weight of the seeds. The M10–M200 treatments increased the FW of the seeds, with the M20 seeds reaching a significantly higher weight. The FW of cotton seed in the M20 treatment increased by 4.81% in comparison with M0 (Fig3).

**Fig 1.**
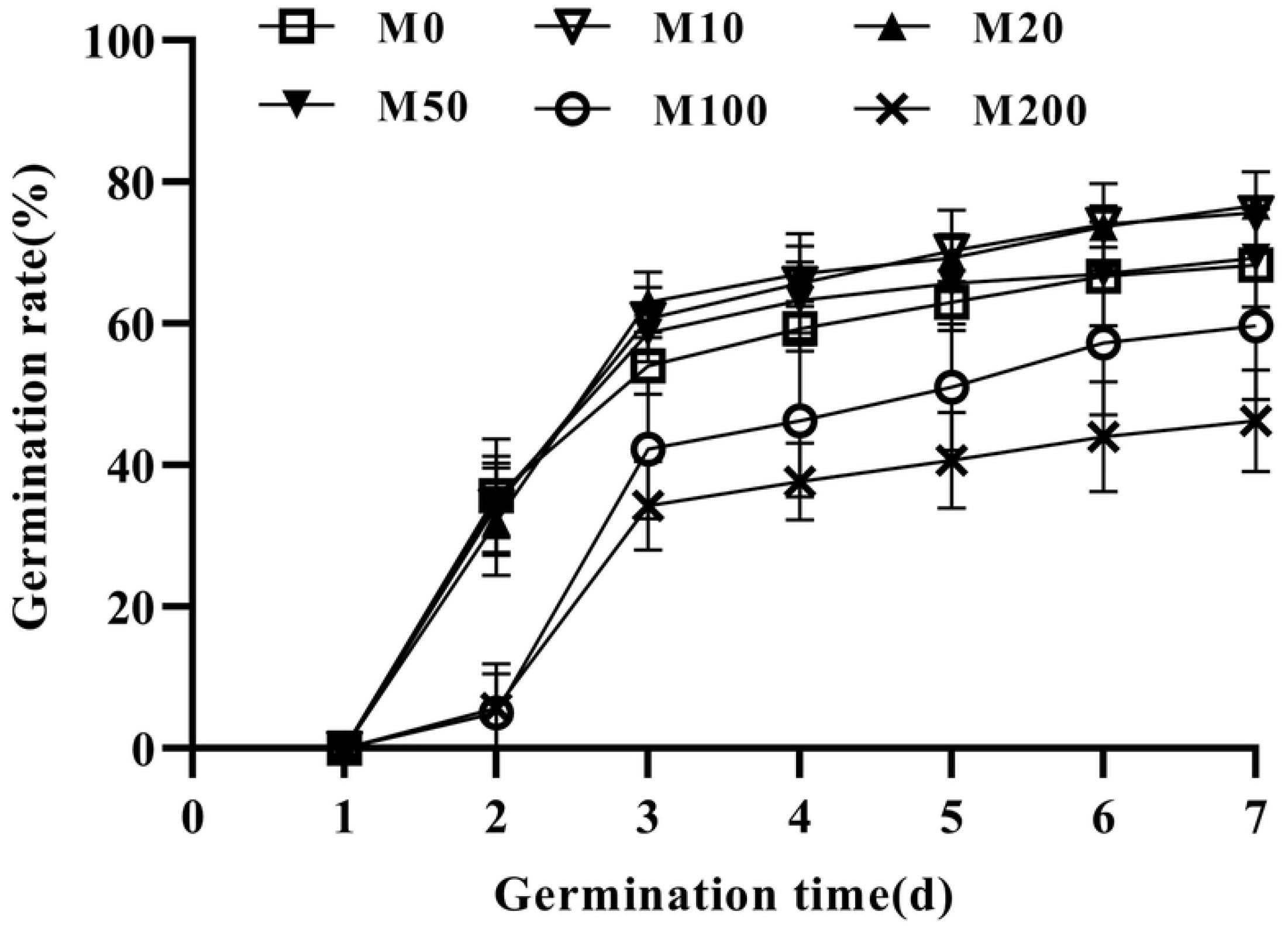
Time-course of changes in germination rate. Treatments were performed with different melatonin concentrations.

**Fig 2.**
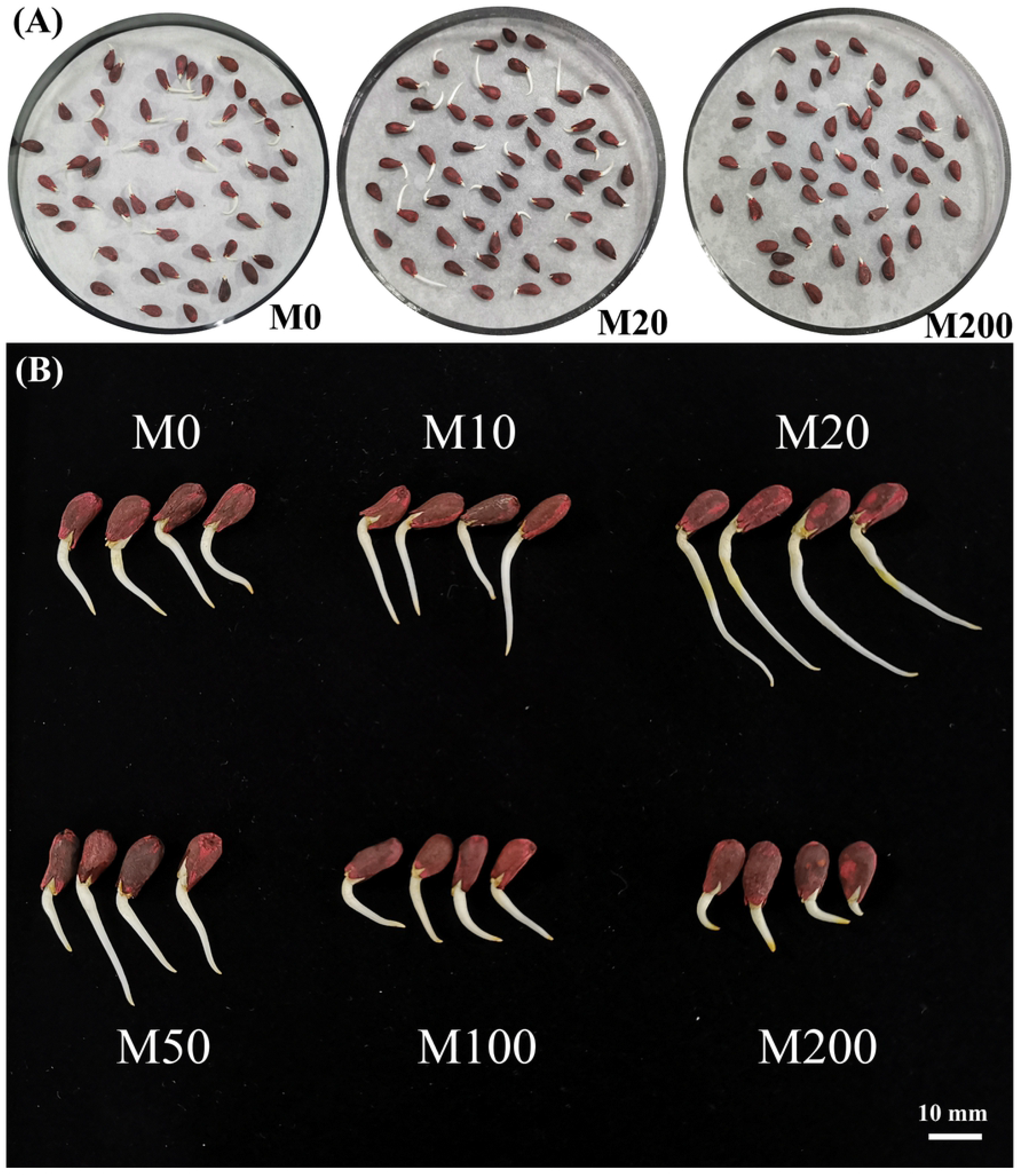
The germination of cotton seeds treated with M0, M20 and M200 after 24 hours of germination (A), and the germination of cotton seeds treated with different concentrations of melatonin on the third day (B)

**Fig 3.**
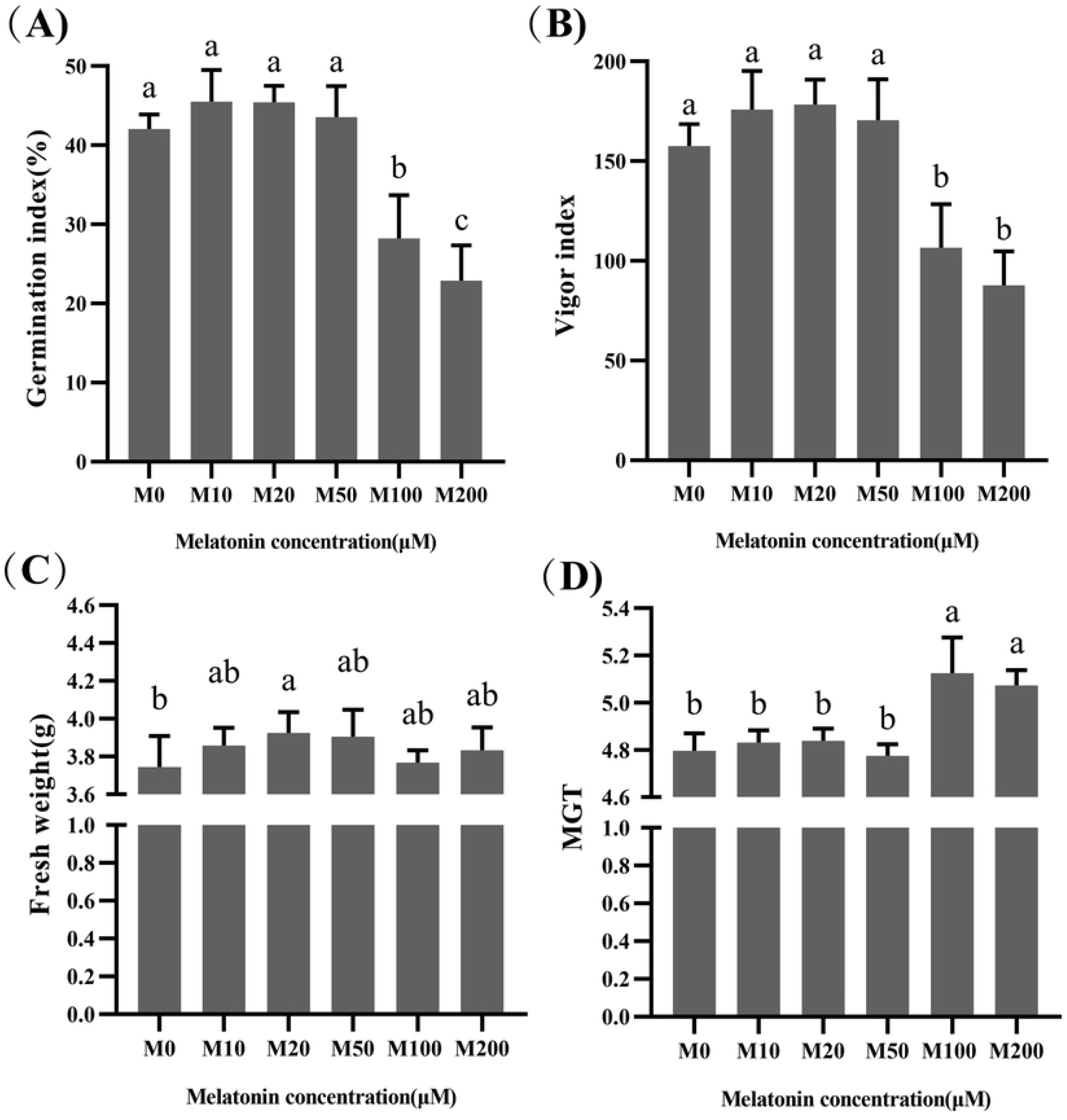
The effects of different concentrations of melatonin on GI (A), VI (B), FW (C) and MGT (D).

### Activity of antioxidant enzymes

Melatonin can scavenge reactive oxygen species by donating electrons and can also affect some oxidative and antioxidant enzymes in cells and tissues through its receptors, for example, by enhancing SOD and POD levels, which act to scavenge free radicals. Among antioxidants, SOD is a particularly active substance in the living body, where it eliminates harmful substances generated by plant metabolism.

In our experiment, SOD and POD activities were determined for each treated seed on days 2, 4, and 6 of germination. Fig 4A shows that the trend in SOD activity was basically uniform across each period, that is, as the concentration of melatonin increased, the content first rose and then decreased. Among the effects of the treatments, that of the M20 treatment was most prominent. On day 2, the activities of SOD enzymes under the M10 and M20 treatments were increased by 28.8% and 39.2%, respectively, and the difference was significant. However, as the melatonin concentration was further increased, SOD activity decreased; when the concentration increased to that of the M200 treatment, SOD activity was reduced by 4.0% compared with the M0 treatment. By day 4 of seed germination, SOD activity did not change, but the activity of each treatment decreased, while that of the M20 treatment was still the highest and significantly different compared to the activity of the M0 treatment. The SOD activity on day 6 was the highest in the three periods measured, while the SOD activity of each treatment tended to be consistent and did not differ, with seeds under the M20 treatment still exhibiting optimal efficacy.

**Fig 4.**
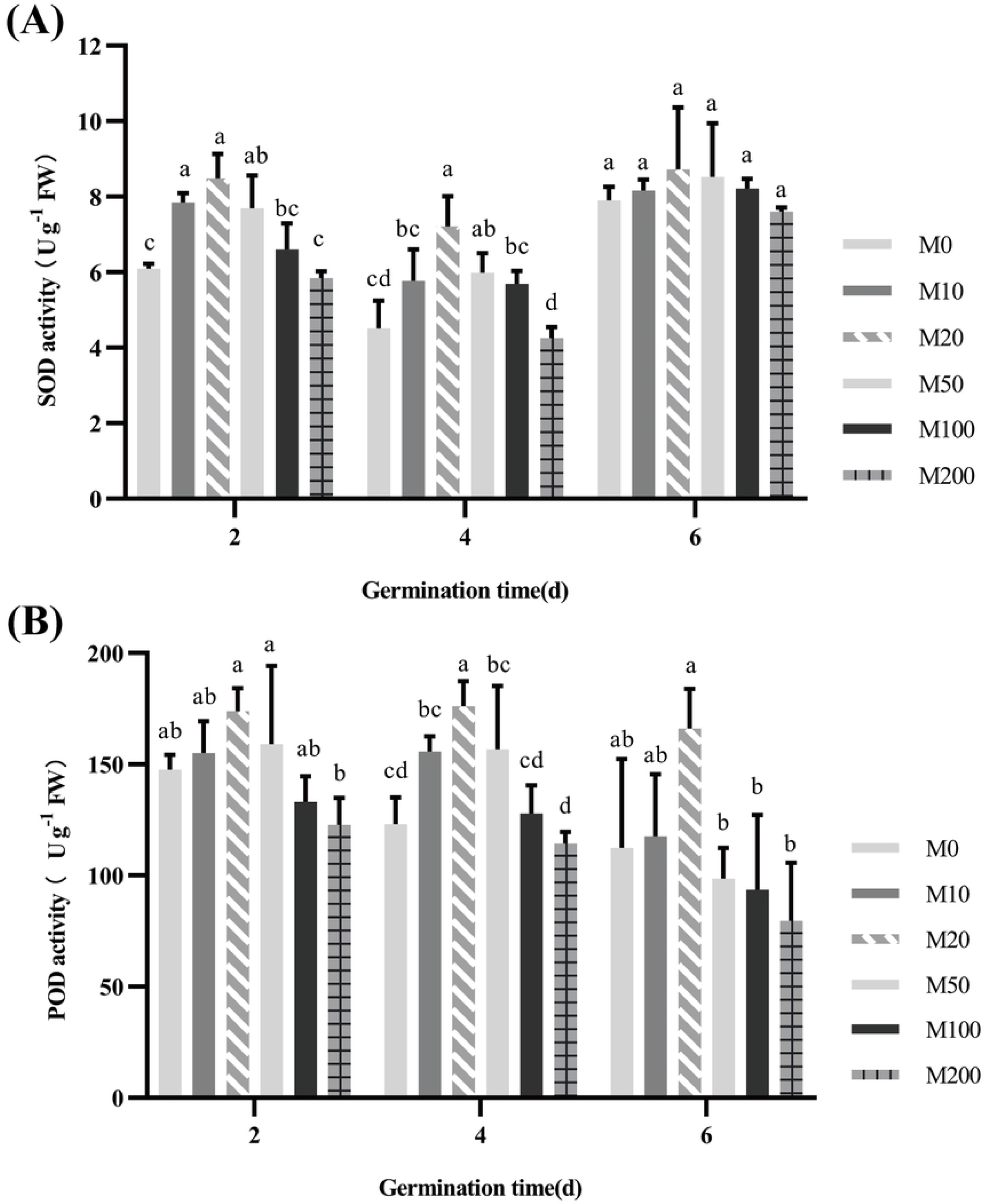
Effect of different concentrations of melatonin on SOD (A) and POD (B).

POD is widely distributed among plants, and its activity reflects the growth and development characteristics of plants. The trend in POD is consistent with that in SOD. By day 2 of seed germination, the POD activities of the M10 and M20 treatment groups continuously increased, by 5.1% and 17.8%, respectively, relative to the M0 treatment, but not significantly. As the melatonin concentration was continually increased, POD activity actually decreased. When the concentration was increased to that of the M200 treatment, POD activity significantly decreased by 16.8% compared with that of the M0 treatment. By day 6, the POD activity of the M20 treatment was still highest, and there was no significant difference compared with that of M0 and M10 treatments. When the melatonin concentration was increased to that of the M50, M100 and M200 treatments, POD activity continued to decrease and reached a significant difference compared with that of the M20 treatment. The POD activity of the M200 treatment group was even less than half of that of the M20 group. Thus, the effect of the M20 treatment on POD was optimal.

### Effects of melatonin on malondialdehyde content

Plant damage is closely related to membrane lipid peroxidation induced by reactive oxygen species accumulation. Malondialdehyde (MDA) is one of the final products of free radical cell membrane lipid peroxidation, and its content can reflect the degree of membrane lipid peroxidation. To assess the oxidative stress generated by different concentrations of melatonin, the extent of membrane lipid peroxidation can be estimated by measuring the MDA content, an indicator of the degree of damage to membrane systems.

Fig 5 shows that the degree of membrane lipid peroxidation of cotton seeds generally increased with time. On days 2, 4 and 6 of seed germination, the MDA content of the M20 treatment was the lowest, but not significantly. Notably, on days 2 and 4, high concentrations of melatonin caused a significant increase in MDA content. Especially on day 2, the MDA content of the M200 treatment reached 2.6 times that of the M0 treatment. At the end of the germination experiment, the MDA content of each treatment tended to be consistent, with significant differences only between the M20 and M50 treatments. Thus, the melatonin concentration of the M20 treatment minimally damaged cell membranes across the whole seed germination trial, while high concentrations of melatonin appeared to cause great damage.

**Fig5.**
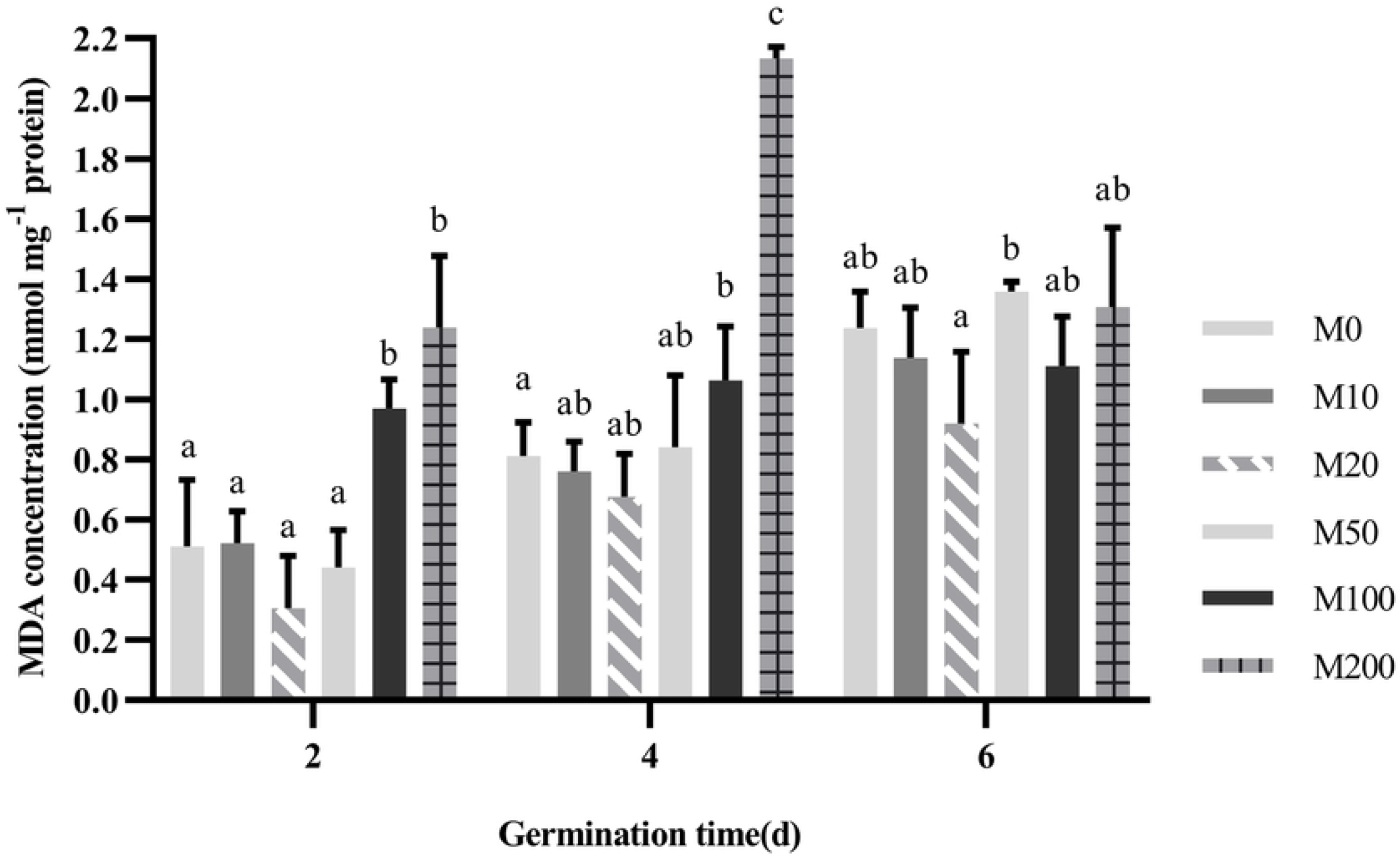
Effect of different concentrations of melatonin on MDA content.

### Effects of melatonin on contents of phytohormones

The phytohormones ABA and GA_3_ are well-known regulators of seed germination. We measured the levels of these phytohormones in seeds subjected to germination trials under different concentrations of melatonin on days 2, 4 and 6 of germination.

Throughout the process of seed germination, GA_3_ content did not continually increase, and it peaked on day 4 of germination. However, by observing the results of germination at each stage separately, we can see that the trends in GA_3_ content were relatively uniform across the different concentrations of melatonin, showing an initial increase followed by a decrease. On day 2 of germination, only the M20 treatment showed a significant increase in GA_3_ content, while other concentrations were associated with different degrees of increase and decrease in GA_3_ content, but without reaching any significant differences. Until day 4 of germination, the GA_3_ content of seeds under the M10 and M20 treatments increased by 79.37% and 151.46% respectively, compared with M0. However, there were no significant differences in GA_3_ content for the other concentration treatments. At the end of the seed germination assay, GA_3_ content decreased as a whole, and the trend among the different concentrations was fairly uniform. GA_3_ content under the M10 and M20 treatments were still highest, while the M100 and M200 treatments exhibited significant decreases in GA_3_ content.

ABA has a dual role of inducing blast cell division to stop as well as initiating and maintaining seed dormancy. As Fig6A shows, the ABA content of seeds gradually increased with days of germination. On day 2, the M20 treatment showed a significant decrease in ABA content, with M100 having the highest ABA content, reaching a significant level. On day 4 of seed germination, seeds under the M20 treatment still performed best. However, the performance of seeds under the M50 and M100 treatments were not the same as on day 2. The ABA content levels under the two treatments were 33.19% and 36.32% lower, respectively, than that under the M0 treatment. There was no significant difference in ABA content between the M100 and M20 treatments, but when the concentration increased to M200, the ABA content increased sharply, through it was not different from that under the M0 treatment. On day 6 of germination, the ABA content was the highest, with only M20 and M50 treatments exhibiting a significant inhibition of ABA content. The M10, M100 and M200 treatments exhibited different degrees of increase or decrease in ABA content, but this did not reach a significant difference compared to M0.

**Fig 6.**
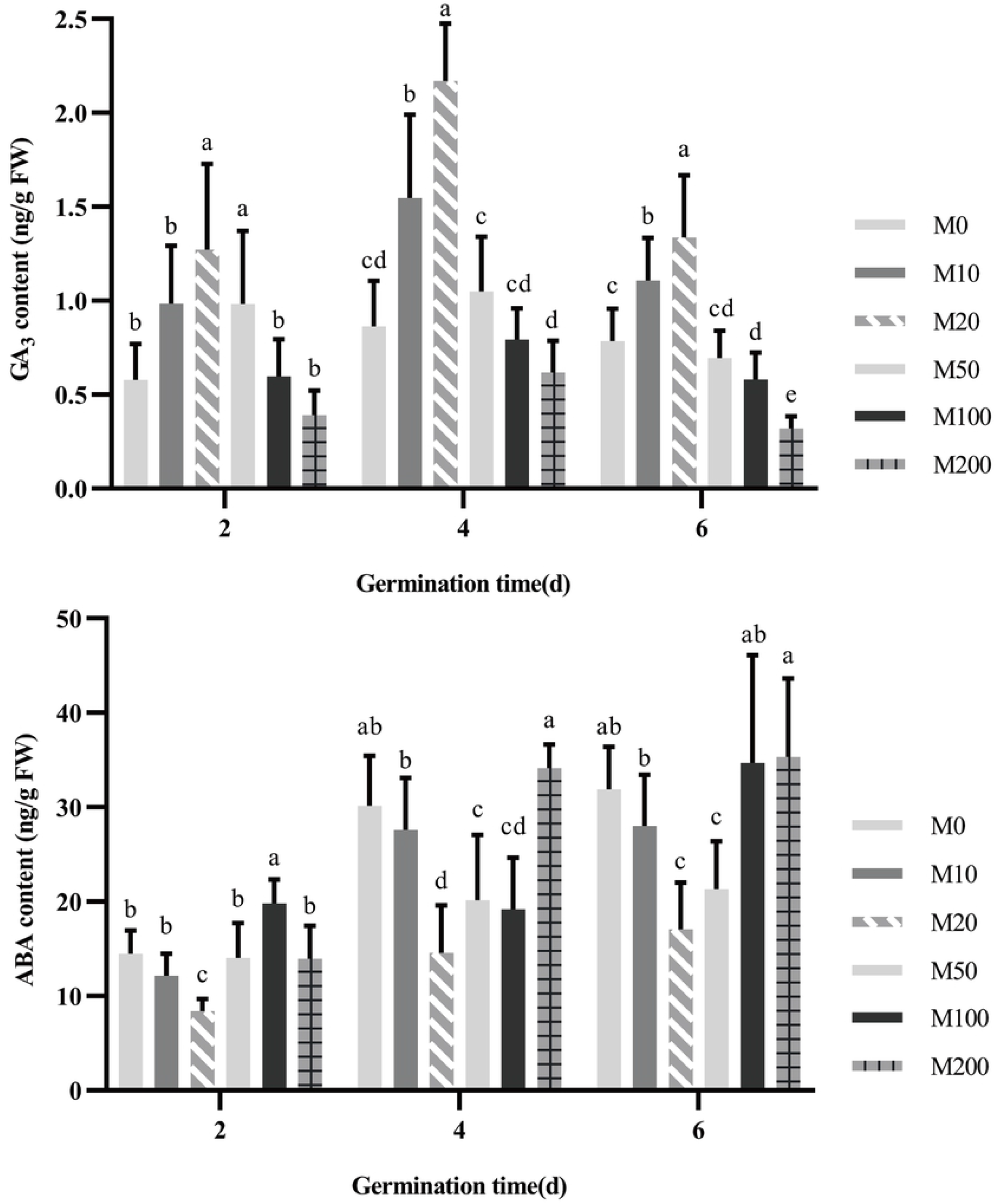
Effects of different concentration of melatonin on contents of GA3(A) and ABA (B).

## Discussion

Melatonin is closely involved in the regulation of development and reproduction [43]. Kolař[44] and Cano[45] confirmed this result in tomatoes and *Chenopodium rubrum*, respectively. In animals, melatonin is not just a hormone. It scavenges a wide range of free radicals in the body, enhances immunity and delays aging [46]. Although melatonin has been widely studied in animals, its possible role in plants is less clear.

Current research indicates that exogenous melatonin has a variety of effects on plants and can be used to regulate plant growth and development. For example, melatonin can promote the growth of lateral roots and slow down the senescence of stems, leaves of plants. In addition, melatonin also improves the resistance of plants to drought, salt damage, high temperatures, cold damage and other adverse conditions, and it can protect cell structural substances and biological macromolecules, such as proteins, from oxidation damage. Melatonin can promote the growth of plants and affect the morphological development of plants. *Hernánde et al. (2004)* found that treatment of etiolated lupins (*Lupinus albus*) with melatonin promoted hypocotyl growth. *Wen et al. (2016)* investigated the effects of melatonin on adventitious root formation of de-rooted tomato seedlings and observed the occurrence of adventitious roots in the seedlings. The induction of adventitious roots in tomato was related to the melatonin concentration of the treatment. The concentration of 50 μmol/L exogenous melatonin was optimal for promoting the emergence of adventitious roots in tomato seedlings.

Seed germination is the process by which the radicle breaks through the seed coat. It is a complex and critical process that determines the establishment of seedlings, and it is also a key period for the establishment of crop populations. Germination includes a range of physical and metabolic events[47]. This stage is greatly affected by the external environment, a period of stress sensitivity and also a critical period for determining the survival of plants under adverse conditions. Seed germination can be considered both the starting point of the plant life cycle and the initial life stage of the plant’s perception of the external environment. Seed germination directly affects the growth and final yield of subsequent cotton seedlings. Therefore, seed germination has important economic and ecological significance *[48, 49]*.

Our experiments assessed GR, GP, GI, VI, MGT and FW in order to accurately evaluate the germination vigor and quality of cotton seeds. GP is used to assess the germination speed and uniformity. The higher the value, the stronger the germination potential is, which is one of the important indicators for assessing seed quality. After the cotton seeds in the experiment were treated with different concentrations of melatonin, the GP of melatonin-treated seeds at concentrations of 10–50μM was improved compared with the control group, with the 20 μM melatonin treatment performing significantly better. In contrast, treatment with 100–200 μM melatonin significantly inhibited GP. Our study revealed that low concentrations (i.e., 10–50 μM) of melatonin could increase the GR of cotton seeds, and 20 μM of melatonin had the strongest effect, but none of them were statistically significant. When the melatonin concentration was increased to 100–200 μM, the GR was significantly inhibited.

GI indicates the germination speed, and the larger the value, the faster the germination speed. The germination test found that treatments with 10–50 μM melatonin promoted the increase of GI. Similarly, M20 still performed best. M100 and M200 significantly reduced GI and reached a highly significant level. However, VI is a better indicator of the subsequent growth of seedlings than is GR. The 10–50 μM melatonin treatments increased the VI compared with the M0 treatment, but not significantly, and the 100–200 μM treatments significantly inhibited the VI. These results indicate that high concentrations of melatonin are likely to have irreversible effects on cotton seeds and even affect the growth of seedlings at later stages. Our test also uses MGT as an indicator to measure seed germination. The lower the MGT, the faster a population of seeds has germinated[50]. The 10–50 μM melatonin treatments have little effect on MGT, and M100 and M200 can significantly increase MGT. This indicator further validates the above conclusions. Our findings are broadly consistent with those of *Simlat et al. (2008)*, who reported that the lowest melatonin concentrations examined (5 and 20 μM) exerted a significantly positive effect on *Stevia* seed germination compared with the 100 and 500 μM treatments. Previous studies have found that melatonin application significantly increased osmopriming effects, such that 50 μM melatonin achieved maximal results[22]. Similar positive effects of melatonin treatment have been reported for red cabbage seeds[23]. However, another study indicated that melatonin had no effect on the germination of cucumber seeds regardless of germination conditions [51].

The germination of the seeds begins with their swelling and water absorption. Accordingly, moisture conditions, a precondition for seed germination, have a significant effect on seed germination. Seeds can only germinate after absorbing a certain amount of water, which is important for the growth of the resulting seedlings. The seeds of different types of plants have different water absorption and germination requirements. Seed absorbs a large amount of water, which also increases dry matter accumulation. FW can directly reflect the water absorption of the seed and the accumulation of biomass. *Bajwa et al. (2014)[52]* reported that the application of low concentrations (10–40 μM) of melatonin increased the FW of *Arabidopsis* compared with high-concentration (200–400 μM) applications. Our results showed that melatonin (10–200 μM) can increase the FW of the seeds, with the 20 μM melatonin treatment having the strongest effect and also reaching a significant level. This result indicates that 20 μM melatonin promotes the water absorption or water absorption rate during seed germination and thus promotes the accumulation of biomass inside seeds, and the inhibitory effect of high concentrations of melatonin on FW was not observed in our study, probably because the highest melatonin concentration examined was insufficient to elicit this response.

As a byproduct of aerobic metabolism[53, 54], reactive oxygen species (ROS) exist in the form of non-free radicals, including hydrogen peroxide (H_2_O_2_), and free radical forms, such as O2•-. The antioxidant system formed during plant evolution contributes to the balance of active oxygen metabolism. The antioxidant defense system includes the enzymatic system and non-enzymatic system. SOD and POD are essential antioxidant enzymes that can effectively eliminate superfluous ROS, such as H_2_O_2_, hydroxyl and superoxide anions, from plant cells[55, 56]. SOD can catalyze the disproportionation of two superoxide anion radicals to form O_2_ and H_2_O_2_, while H_2_O_2_ is further decomposed by other antioxidant enzymes such as CAT. As melatonin and its metabolites are known as endogenous free radical scavengers, broad-spectrum antioxidants and H_2_O_2_ scavengers, it is important to determine whether melatonin can also reduce the inhibition of seed germination by ROS [42, 57]. Our study found that different concentrations of melatonin have effects that include low concentration promotion and high concentration inhibition of SOD and POD activity during seed germination, and the peak activity of both occurred under the 20 μM treatment. This finding suggests that low concentrations of melatonin increase the activity of antioxidant enzymes including SOD and POD, and exogenous applications of melatonin appear to play a role in the first line of defense against oxidative stress. However, there are studies that offer the opposite conclusion, *Shi et al. (2015a)* [58] and *Shi et al. (2015b)[59]* found no significant effects of melatonin on antioxidant enzyme activity under controlled conditions. Other researchers have found that melatonin not only directly eliminates ROS, but also upregulates the activity of various antioxidant enzymes[60]. This was in accord with observations in the current study.

The accumulation of ROS and free radicals in organisms often leads to membrane lipid peroxidation and causes metabolic disorders. The integrity of cell structure is the basis of seed vigor, so once the cell membrane system is destroyed, leakage increases throughout the progression of germination. MDA is a product of lipid peroxidation, so it represents the level of lipid peroxidation and the degree of membrane damage in cells. The MDA content under the 20 μM melatonin treatment was the lowest, but not significantly, while the high-concentration (100– 200 μM) treatments significantly increased the MDA content. This indicates that low concentrations of melatonin can alleviate peroxidative stress during seed germination but that it can still have a negative effect at high concentrations.

Seed germination, as the beginning of the agricultural cycle from the perspective of the plant, is the first stage of crop growth. Hormones inside the seed control the germination of seeds. For example, ethylene and gibberellin weaken the seed coat and break seed dormancy to promote seed germination. Numerous studies have shown that melatonin has synergistic and antagonistic relationships with various other plant hormones, for example, promoting plant root and young leaf growth at low concentrations, which is similar in action to auxin (IAA).

There are few studies on the effects of melatonin treatment on gibberellin content. Under salt stress, melatonin treatment of cucumber seedlings can upregulate the expression of GA biosynthesis genes and increase the content of active GAs such as GA_3_ and GA_4_, thus promoting subsequent germination processes under salt stress[61, 62]. The present study shows that although changes in GA_3_ content may slightly differ among the days of seed germination, both 10 μM and 20 μM melatonin treatments, especially the latter, can significantly increase the GA content of seeds. Compared with the control treatment, melatonin had no effect on GA_3_ content and even inhibited it at concentrations of 50–200 μM. At present, there are few studies on the effects of melatonin treatment on gibberellin content. However, melatonin treatment has been reported to increase the content of active GAs such as GA_3_ and GA_4_ in cucumber seedlings, thus promoting the germination process under salt stress inhibition[61, 63].

ABA is the most important plant hormone regulating seed dormancy and germination. It induces and maintains seed dormancy, inhibits seed germination and regulates seedling growth and development. Melatonin treatment can induce the decreases in ABA level during seed germination under salt stress by inducing upregulation of ABA catabolic genes and downregulation of ABA biosynthesis genes[61]. In the heat-induced senescence of the perennial ryegrass (*Lolium perenne* L.) leaves, exogenous melatonin treatment reduced ABA content and delayed its aging process[64]. In our study we found that low concentrations of melatonin inhibited ABA content, while 100–200 μM melatonin treatments promoted ABA content in seeds, but not significantly.

## Conclusions

The present study demonstrates the positive effects of melatonin on seed germination in cotton. Low concentrations of melatonin had a positive effect on cotton seed germination compared with high concentrations, of which 20μM melatonin was the optimal treatment. This concentration also resulted in low levels of MDA concentration as well as high SOD and POD activity. Additionally, melatonin may promote seed germination by regulating the endogenous synthesis of phytohormones, as the hormone content of seeds treated with low concentrations of melatonin, which promoted seed germination, also showed a corresponding pattern, in which the 20μM melatonin treatment achieved the best germination and significantly increased GA_3_ content and reduced ABA content. Correspondingly, high concentrations of melatonin generally showed no effect or even inhibited germination somewhat.

## Abbreviations

GR: germination rate
GP: germination potential
GI: germination index
VI: vigor index
MGT: mean germination time
FW: fresh weight
ROS: reactive oxygen species
SOD: superoxide dismutase
POD: peroxidase
MDA: malondialdehyde
ABA: abscisic acid
GA: gibberellin

## Acknowledgements

The authors are grateful to the anonymous reviewers for their valuable comments and suggestions. This work was supported by the National Natural Science Foundation of China (No. 31871569, 31171495, 31301270 and 31571610).

